# Intramolecular domain dynamics regulate synaptic MAGUK protein interactions

**DOI:** 10.1101/462358

**Authors:** Nils Rademacher, Benno Kuropka, Stella-Amrei Kunde, Markus C. Wahl, Christian Freund, Sarah A. Shoichet

## Abstract

PSD-95 MAGUK family scaffold proteins are multi-domain organisers of synaptic transmission that contain three PDZ domains followed by an SH3-GK domain tandem. This domain architecture allows coordinated assembly of protein complexes composed of neurotransmitter receptors, synaptic adhesion molecules, cytoskeletal proteins and downstream signalling effectors. Here we show that binding of monomeric PDZ_3_ ligands to the third PDZ domain of PSD-95 induces functional changes in the intramolecular SH3-GK domain assembly that influence subsequent homotypic and heterotypic complex formation. We identify PSD-95 interactors that differentially bind to the SH3-GK domain tandem depending on its conformational state. Among these interactors we further establish the heterotrimeric G protein subunit Gnb5 as a PSD-95 complex partner at dendritic spines. The PSD-95 GK domain binds to Gnb5 and this interaction is triggered by PDZ_3_ ligands binding to the third PDZ domain of PSD-95, unraveling a hierarchical binding mechanism of PSD-95 complex formation.

## Introduction

Excitatory synapses are the contact sites through which neurons communicate which each other. These synapses are asymmetric structures that are formed by pre- and postsynaptic terminals containing distinct sets of proteins. Incoming action potentials are converted into chemical signals (neurotransmitters) at presynaptic terminals, which subsequently pass through the synaptic cleft and are reconverted into electrical signals at postsynaptic sites (Lisman et al., 2007). These synaptic contacts are not static but are able to undergo structural changes and thereby modify neuronal network computation (Nishiyama and Yasuda, 2015). At postsynaptic sites, interacting proteins are densely packed into a sub-membrane structure called the postsynaptic density (PSD) (Sheng and Hoogenraad, 2007). Scaffold proteins of the PSD-95 family membrane-associated guanylate kinases (MAGUKs) are highly abundant components of the PSD and function as central regulators of postsynaptic organisation (Zhu et al., 2016a). PSD-95 family MAGUKs contain three PDZ domains that are known to directly interact with N-methyl-D-aspartate (NMDA) and α-amino-3-hydroxy-5-methyl-4- isoxazolepropionic acid (AMPA) receptor C-termini (Kornau et al., 1995; Leonard et al., 1998; Dakoji et al., 2003) followed by an SH3 - guanylate kinase (GK) domain tandem (Funke et al., 2005). The MAGUK SH3 domain lost its function to bind proline-rich peptides; instead it forms an intramolecular interaction with the GK domain (McGee et al., 2001). Similarly, the PSD-95 GK domain is atypical in that it is unable to phosphorylate GMP but has evolved as a protein interaction domain (Johnston et al., 2011). Until now, binding of known interactors to the GK domain typically involves residues of the canonical GMP-binding region (Reese et al., 2007; Zhu et al., 2011; Zhu et al., 2016b). This modular array of protein interaction domains allows PSD-95 MAGUKs to function as bidirectional organisers of synaptic function. First, neurotransmitter receptors can be incorporated or removed from postsynaptic membranes, depending on molecular interactions with these sub-membrane scaffold proteins. Second, together with other scaffold proteins at postsynaptic sites, they align downstream effectors and cytoskeletal proteins. Accordingly, PSD-95 family MAGUKs are essential for the establishment of long-term potentiation (LTP) by regulating the content of AMPA receptors at dendritic spines (Ehrlich and Malinow, 2004; Opazo et al., 2012; Sheng et al., 2018). In line with this is the observation that acute knockdown of PSD-95 MAGUKs leads to a decrease in postsynaptic AMPA and NMDA receptor-mediated synaptic transmission as well as a reduction in PSD size (Chen et al., 2015). Taken together, exploring protein complex formation directed by PSD-95 MAGUK family members is of central interest to understand synaptic regulation. We have previously shown that synaptic MAGUK proteins oligomerise upon binding of monomeric PDZ_3_ ligands (ligands that specifically bind to the third PDZ domain) (Rademacher et al., 2013) and speculated that ligand - PDZ_3_ domain binding induces conformational changes in the C-terminal domains that lead to complex formation. Our initial observations of PDZ ligand-induced effects in MAGUK proteins have been recently supported by other studies (Zeng et al., 2016; Zeng et al., 2017).

In this study, we use a bimolecular fluorescence complementation (BiFC) assay to show that PSD-95 oligomerisation is triggered by PDZ_3_ ligands and dependent on the C-terminal SH3-GK domain tandem. Moreover, we identify synaptic interaction partners of PSD-95 C-terminal domains by quantitative mass spectrometry and provide evidence that the heterotrimeric G protein subunit Gnb5 is a novel GK domain interactor and that its ability to bind to PSD-95 is likewise promoted by ligand binding to the PSD-95 PDZ_3_ domain.

## Results

### Ligand binding to PSD-95 PDZ_3_ domains facilitates oligomerisation guided by its C-terminal module

We are interested in the functional coupling of PDZ_3_ domains with the adjacent SH3-GK domain tandem in the synaptic scaffold protein PSD-95 (PSG module, see **Figure 1A** for domain structure) and the relevance of ligand - PDZ_3_ domain interactions on PSD-95 complex formation. To explore this idea, we built on our previous work with tagged cytosolic PDZ3 ligands (Rademacher et al., 2013) and we have now designed a cell-based assay to directly monitor the proximity of PSD-95 molecules by bimolecular fluorescence complementation (BiFC). Expression constructs of PSD-95 were fused to non-fluorescent halves of EYFP (N-terminal half = YN and C-terminal half = YC) and coexpressed with the established PDZ domain ligand Neuroligin-1 (NLGN1) in HEK cells. NLGN1 is a synaptic adhesion molecule that specifically binds to the third PDZ domain of PSD-95 (Irie et al., 1997). Coexpression of the per se non-fluorescent PSD-95-YN and PSD-95-YC constructs (together referred to as WT/WTsplitEYFP) with full-length NLGN1 led to the formation of fluorescent PSD-95 complexes that were located at the cell membrane, recapitulating the natural localisation of the endogenous protein complexes (**Figure 1B**). Next, we quantified the formation of fluorescent complexes by flow cytometry (**Figure 1C**). Interestingly, upon coexpression of mutant NLGN1 constructs that carry two alanine substitutions within the C-terminal PDZ_3_ ligand sequence (mutNLGN1: C-terminus TTRV ▸ T**A**R**A**), the detected fluorescence intensity decreased by approximately 40% (**Figure 1C**). Fluorescent signals were nearly undetectable following coexpression of PSD-95-YC with the scaffold-incompetent PSD-95 point mutant PSD-95-YN L460P, together with either NLGN1 or mutNLGN1 (**Figure 1C**). Leucine 460 is an internal SH3 domain residue and the L460P mutation has been shown to specifically disrupt the intramolecular SH3-GK domain interaction (**Supplemental Figure 1**) (McGee and Bredt, 1999; Shin et al., 2000) that is one of the hallmark features of MAGUK proteins (Tavares et al., 2001). Interestingly, this amino acid exchange does not interfere with PDZ_3_ ligand binding (Rademacher et al., 2013) but strongly abolishes PSD-95 complex assembly (**Figure 1C**). We assume that in the context of the full-length protein, the L460P mutation likewise weakens the (intramolecular) interaction between the SH3 and GK domain, which would then result in a constitutively ‘open’ conformation. This profound negative effect that we observe following a targeted amino acid exchange in the SH3 domain highlights the importance of the SH3-GK domain tandem for its involvement in regulated PSD-95 oligomerisation.

**Figure 1.**
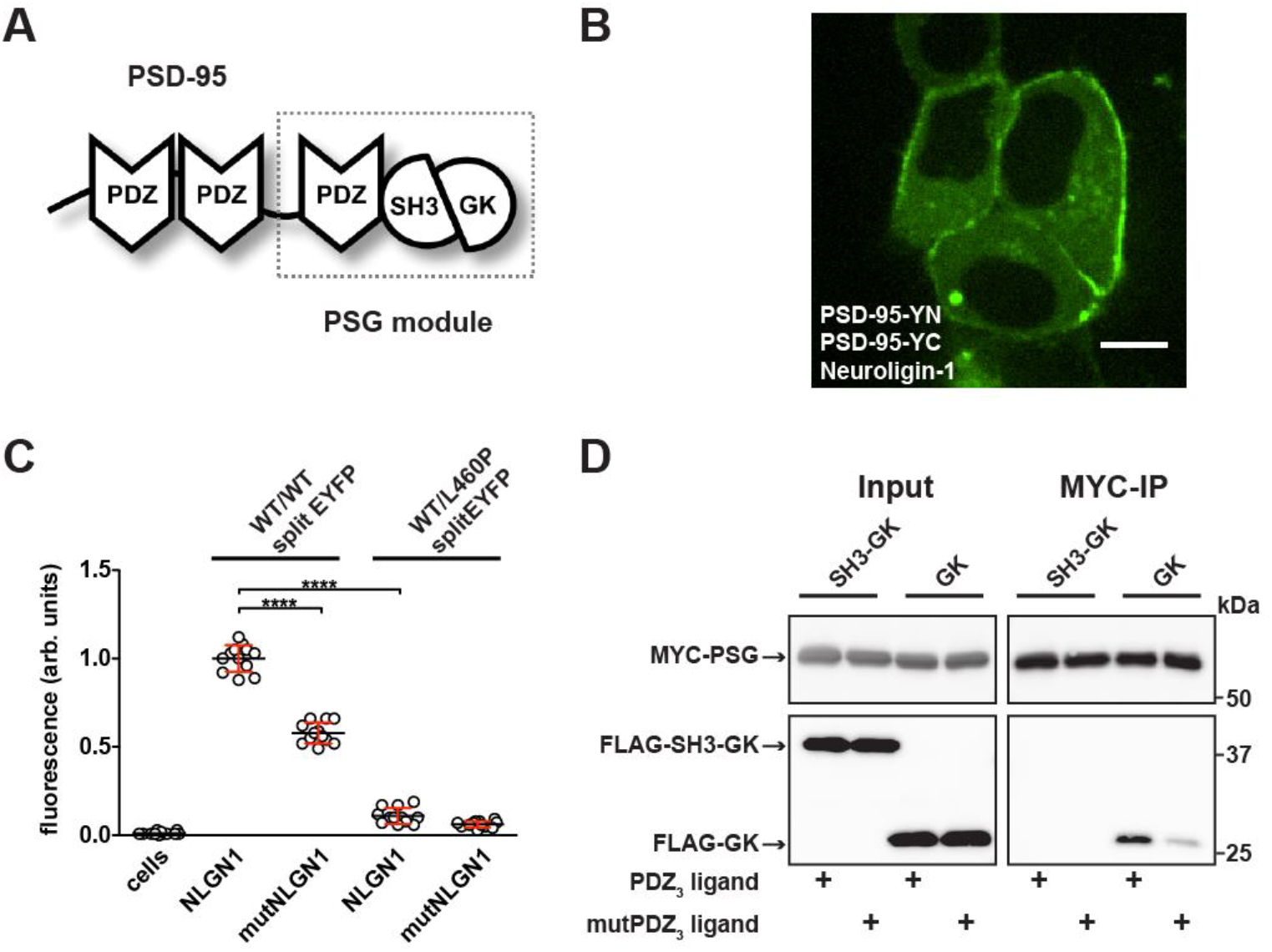
PDZ_3_ ligand-induced dynamics in the PDZ_3_-SH3-GK module facilitate oligomerisation. **A)** Schematic representation of the PSD-95 domain organisation. PSD-95 contains three PDZ domains followed by a SH3-GK domain tandem. The PSG module (**P**DZ_3_-**S**H3-**G**K) is common to the MAGUK protein family. **B)** Live-cell microscopy of HEK-293T cells transfected with PSD-95-YN, PSD-95-YC and NLGN1 reveals a membrane associated localisation of the refolded complex (transfection corresponding to WT/WTsplitEYFP plus NLGN1 in Figure 1C). Scale bar: 10 μm. **C)** PSD-95 oligomerisation assay based on BiFC. HEK-293T cells were triple-transfected with the displayed DNA constructs and EYFP refolding was assessed by flow cytometry. Formation of oligomeric fluorescent complexes is effective in the presence of wild-type Neuroligin-1 (NLGN1). Fluorescence is reduced by either site-directed mutagenesis of the NLGN1 PDZ_3_ ligand C-terminus (mutNLGN1: TTRV ▸ T**A**R**A**), or a targeted amino acid exchange in the PSD-95 SH3 domain (L460P). The dot plot indicates mean values (black horizontal bar) with SD (red vertical bar), based on twelve individual measurements (dots) that originate from four independent experiments (results from each experiment are triplicates for each DNA construct combination). Data was analysed by one-way ANOVA / Sidak’s multiple comparisons test. ****p < 0.0001. **D)** MYC-PSG and FLAG-SH3-GK or FLAG-GK were coexpressed together with either PDZ_3_ ligand or mutPDZ_3_ ligand constructs. Upon MYC-PSG IP, proteins were analysed by western blot with αFLAG antibodies. Coexpression of the PDZ_3_ ligand enhanced the coIP of PSG and GK, whereas coIP of PSG and SH3-GK was negligible regardless of whether or not the PDZ_3_ ligand construct was coexpressed. The following source data is available for this figure: **Source data 1.** Source data for ***Figure 1C***.

### Ligand binding to PSD-95 PDZ_3_ facilitates an ‘open’ SH3-GK state that frees both domains for binding in *trans*

In line with our BiFC assay results, we have previously reported that PSD-95 constructs (full-length and PSG module), efficiently oligomerise and coprecipitate upon binding of a PDZ_3_ ligand (Rademacher et al., 2013). Moreover, the observation by NMR spectroscopy that the PSG module forms a dynamic modular entity (Zhang et al., 2013) led us to hypothesise that ligand binding to PDZ_3_ might influence intramolecular SH3-GK domain assembly, facilitating the formation of domain swapped oligomers (McGee et al., 2001; Ye et al., 2018). Specifically, we asked whether the ligand - PDZ_3_ domain interaction might release the intramolecular SH3-GK domain assembly, thereby allowing other domains and proteins to interact in *trans*. To explore this idea, we assessed which PSD-95 domains are able to interact in *trans* upon PDZ_3_ ligand binding to proteins that harbour the PSG module, using a coimmunoprecipitation experiment designed accordingly. We expressed the PSG module together with PDZ_3_ ligand constructs consisting of the last 10 amino acids (DTKNYKQTSV) of the established PDZ_3_ binder CRIPT (Niethammer et al., 1998) fused to the monomeric red fluorescent protein mCherry (referred to as ‘PDZ_3_ ligand’). As a control, we coexpressed similar constructs carrying two amino acid exchanges within the PDZ_3_ ligand sequence (DTKNYKQ**A**S**A**, referred to as ‘mutPDZ_3_ ligand’). Upon triple transfection with either a GK or an SH3-GK domain construct, the PSG modules were precipitated, and copurified proteins were analysed by western blot (**Figure 1D**). The SH3-GK construct did not coprecipitate with the PSG module regardless of whether it was coexpressed with wild-type or mutant PDZ_3_ ligands. This may be due to a constitutive intramolecular association of the SH3 and GK domains, leading to a ‘closed’ SH3-GK assembly, with no ability to bind a PSG module in *trans*. The GK domain alone, however, coprecipitated effectively with the PSG module, when expressed in the presence of functional PDZ_3_ ligands. These data suggest that binding of a PDZ_3_ ligand renders the PSG module ‘interaction-competent’, *i.e.* it facilitates formation of a conformational state in which it is able to bind isolated GK domain constructs in *trans*. In this experiment, the intramolecular SH3-GK domain assembly resembles the ‘interaction-incompetent’ state, and the SH3 domain autoinhibits the GK domain’s interaction activity.

### The SH3-GK assembly state influences PSD-95 interactions with synaptic proteins

Based on the above results, we propose that PSD-95 C-termini can adopt different functional states depending on whether or not PDZ_3_ ligands are bound to PSD-95 PDZ_3_ domains, *i.e.* that ligand binding induces a loosening of the intramolecular SH3-GK domain assembly and renders the SH3-GK domain tandem ‘interaction-competent’. In order to identify interactors that differentially bind to PSD-95 C-termini in an ‘open state’ versus PSD-95 molecules where the GK domain is autoinhibited by an intramolecular interaction with the adjacent SH3 domain, we utilised a quantitative proteomics strategy. In a reductionist approach, we mimic the open and closed states with different bacterially expressed GST fusion proteins: a GST-GK construct serves as the ‘interaction-competent’ GK domain state, whereas a GST-SH3-GK domain fusion protein reflects the autoinhibited domain assembly. By performing GST pull-downs from crude synaptosome preparations followed by quantitative mass spectrometric analysis we aimed to identify novel proteins that preferentially bind to the ‘open’ or ‘closed’ state of the PSD-95 C-terminal domains (**Figure 2A**).

**Figure 2.**
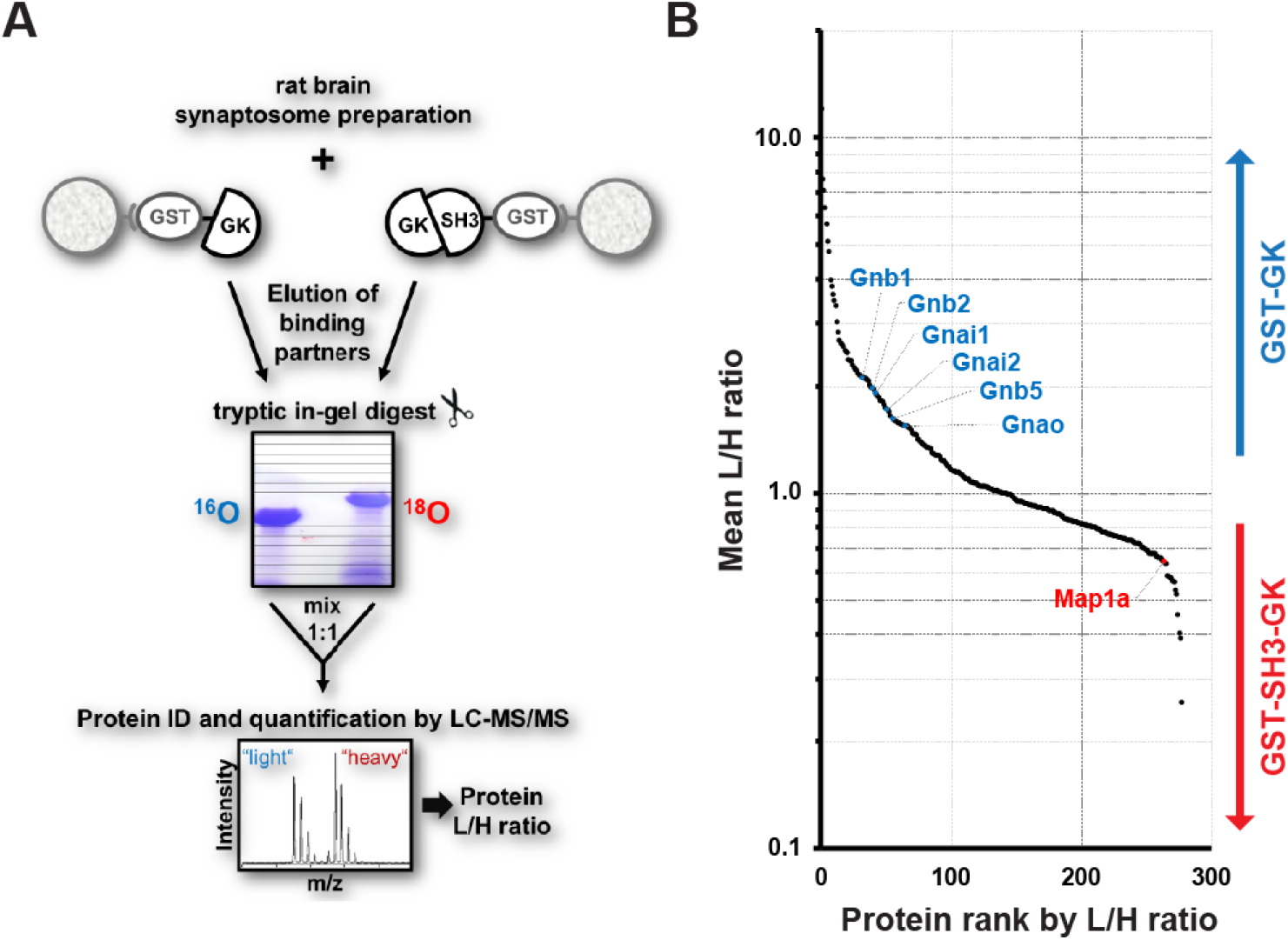
Identification of interactors that differentially bind to PSD-95 C-terminal domains. **A)** Schematic representation of the quantitative mass spectrometry experiment to identify PSD-95 GK domain interactors from rat synaptosomes by GST pull-down of bacterially expressed GST-GK or GST-SH3-GK constructs and ^18^O-labeling. **B)** GST pull-downs were performed in triplicates and 278 interacting proteins that passed our threshold settings were identified and quantified by mass spectrometry. Proteins are ranked by their mean L/H ratio indicating preferential enrichment with either GST-GK or GST-SH3-GK constructs. The heterotrimeric G protein subunit Gnb5 was found to be enriched in the GST-GK fraction relative to the GST-SH3 GK fraction and selected for further studies. The following figure supplement is available: **Supplemental Table 1.** Source data and supplement for ***Figure 2B***.

Bacterially expressed GST-GK vs. GST-SH3-GK constructs were incubated with crude synaptosome preparations of whole rat brains in triplicates. Interacting proteins were eluted from the beads and separated by SDS-PAGE. Enzymatic ^16^O/^18^O-labelling during tryptic in-gel digestion was used for relative quantification of proteins by nanoLC-MS/MS analysis. Proteins enriched by GST-GK were labelled light (^16^O), while proteins enriched by GST-SH3-GK were labeled heavy (^18^O). In total, we reproducibly identified and quantified 278 proteins (**Supplemental Table 1**). Remarkably, 208 (≈ 75%) of these have been recently reported to be present in postsynaptic density fractions isolated from prefrontal cortex (Wilkinson et al., 2017). Moreover, we also identified the known GK-domain interacting proteins Map1A (Reese et al., 2007), Mark2 (Wu et al., 2012), Dlgap2 (Takeuchi et al., 1997) and Srcin1/p140CAP (Fossati et al., 2015), validating the general success of our approach. Potential binders to the GST-GK construct are expected to be enriched in their light form (L/H ratio > 1), while binders to the GST-SH3-GK construct are expected to be enriched in their heavy form (L/H ratio <1). Unexpectedly, we isolated several heterotrimeric G protein subunits enriched in the protein fractions that bind preferentially to the GST-GK construct (**Figure 2B**). Of special interest was the guanine nucleotide binding protein beta 5 (Gnb5), which is a signalling effector downstream of GPCRs that exhibits inhibitory activity in neurons (Xie et al., 2010; Ostrovskaya et al., 2014). Gnb5 contains an N-terminal α-helix followed by a β-sheet propeller composed of seven WD-40 repeats (Cheever et al., 2008). Gnb5 is specifically expressed in brain (Watson et al., 1994) and mutations in the Gnb5 gene cause a multisystem syndrome with intellectual disability in patients (Lodder et al., 2016).

### Gnb5 is a novel synaptic PSD-95 complex partner

In order to verify Gnb5 as a potential binding partner from the above mass spectrometry result, we performed a GST pull-down from crude rat brain synaptosomes and analysed the associated proteins by western blot (**Figure 3A**). We could not detect Gnb5 in the bead control pull-down lane and almost no Gnb5 was detectable in the GST-SH3-GK lane. However, a clear Gnb5 signal was present in the GST-GK lane, supporting our quantitative mass spectrometry results and suggesting that a Gnb5 - GK domain interaction is favoured over a Gnb5 - SH3-GK domain interaction. Additionally, we observed a preferred interaction of overexpressed Gnb5 with the isolated GK domain compared to SH3-GK domain constructs in COS-7 cells. Upon IP of Gnb5 tagged with the green fluorescent protein Clover (Lam et al., 2012) with αGFP antibodies, the GK domain construct coprecipitates far more efficiently than does the SH3-GK domain (**Figure 3B**).

**Figure 3.**
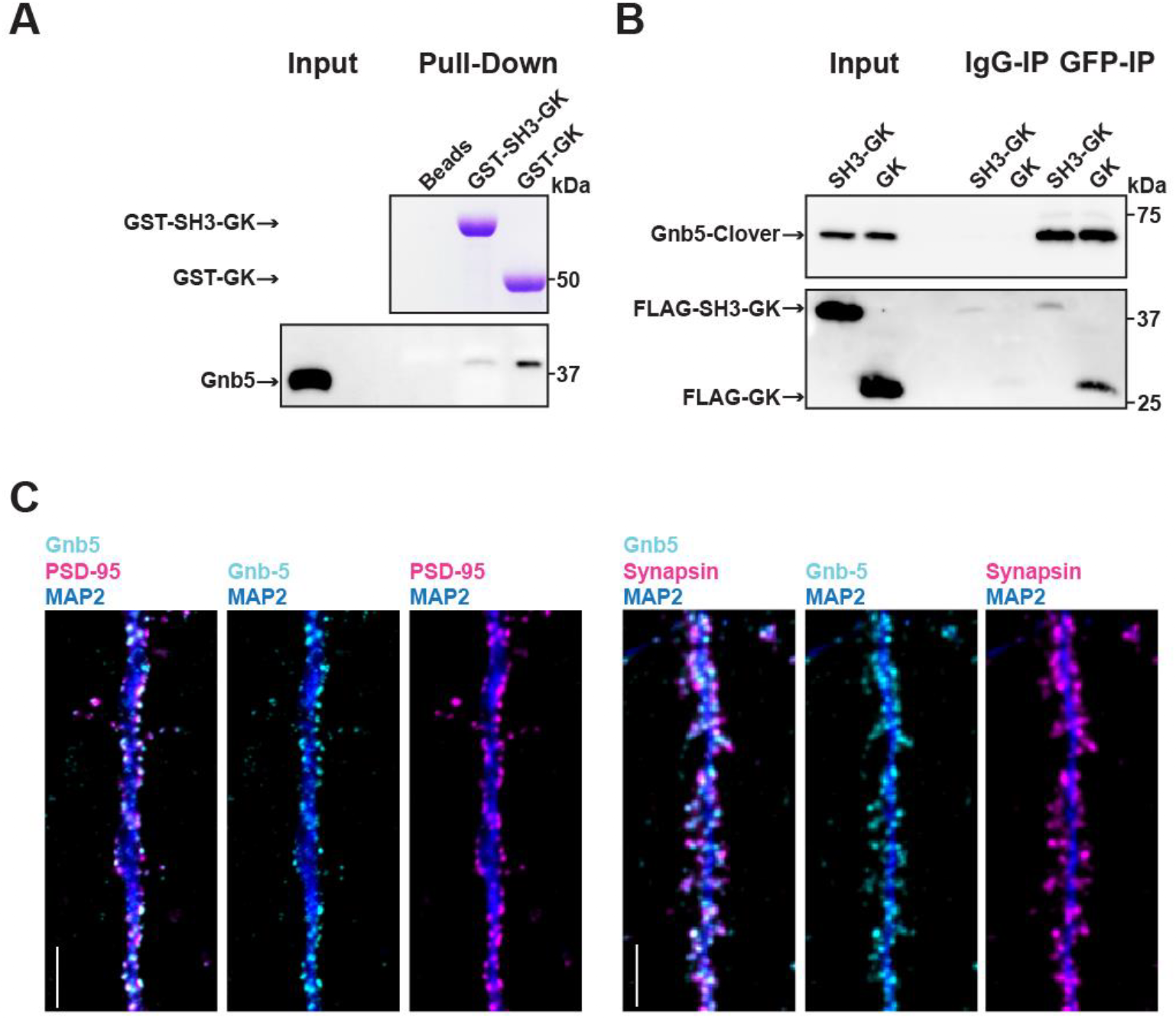
The heterotrimeric G protein subunit Gnb5 is a novel PSD-95 interactor. **A)** GST pull-down from crude synaptosomal proteins (comparable amounts of GST tagged proteins observable by Coomassie, upper panel) enabled comparison of Gnb5 binding to the GK domain alone versus the SH3-GK domain. Gnb5 is effectively enriched in the GST-GK pull-down compared to bead controls or GST-SH3-GK pull-downs, as observed by western blot with a commercially available αGnb5 antibody (lower panel). **B)** CoIP experiment of tagged Gnb5 (Gnb5-Clover) with tagged SH3-GK or GK (FLAG-SH3-GK or FLAG-GK). Immunoprecipitation of Gnb5-Clover with αGFP antibody efficiently copurified the GK-domain construct (observed via western blot with αFLAG antibodies, lower panel). **C)** Cultures of rat hippocampal neurons (E18) were fixed at DIV21 and stained for Gnb5 together with the dendritic marker MAP2 (microtubule-associated protein 2) and either the postsynaptic protein PSD-95 (left panel) or the presynaptic marker Synapsin (right panel) and respective fluorescent secondary antibodies, and visualised by confocal microscopy. Scale bars: 5 μm.

Our *in vitro* experiments clearly indicate that Gnb5 is an interactor of PSD-95 C-terminal domains. However, our interaction data do not clearly indicate in which subcellular compartment Gnb5 and the Gnb5 - PSD-95 complex is located. To explore this, we immunostained cultures of dissociated cells from rat hippocampi and analysed the subcellular distribution of endogenous proteins. We stained fixed cultures (DIV21) with antibodies against Gnb5 and costained for the dendritic marker MAP2 and PSD-95. Gnb5 staining was present in neuronal dendrites, where the signal overlaps with the PSD-95 staining (**Figure 3C**). Additionally, we stained neurons with antibodies against Gnb5, MAP2 and the presynaptic marker Synapsin. In these experiments, the Gnb5 signal is adjacent to the presynaptic Synapsin signal (**Figure 3C**). Together, these findings strongly support the idea that Gnb5 and PSD-95 are protein complex partners at postsynaptic sites of hippocampal neurons.

### Regulation of PSD-95 complex formation

Our data indicate that Gnb5 interacts differentially with PSD-95 C-terminal constructs and we observe that PSD-95 and Gnb5 exhibit overlapping expression exclusively at postsynaptic sites. We next set out to determine if the PSD-95 - Gnb5 interaction is indeed influenced by the presence of synaptic PDZ_3_ ligands, as we initially hypothesised. We coexpressed PSD-95 with PDZ_3_ ligand constructs as in previous experiments, together with Gnb5. Following IP of PSD-95, the precipitates were analysed by western blot: the presence of PDZ_3_ ligands indeed triggered coimmunoprecipitation of Gnb5 and PSD-95, which supports the idea that ligand binding to PDZ_3_ indirectly affects protein-protein interactions at neighbouring domains. Gnb5 lacking the N-terminal α-helix (shortGnb5) coprecipitated somewhat less efficiently than the full-length protein (**Figure 4A**), suggesting that this N-terminal region of Gnb5 (amino acids 1-33) is important for the PDZ_3_ ligand-mediated interaction with PSD-95.

**Figure 4.**
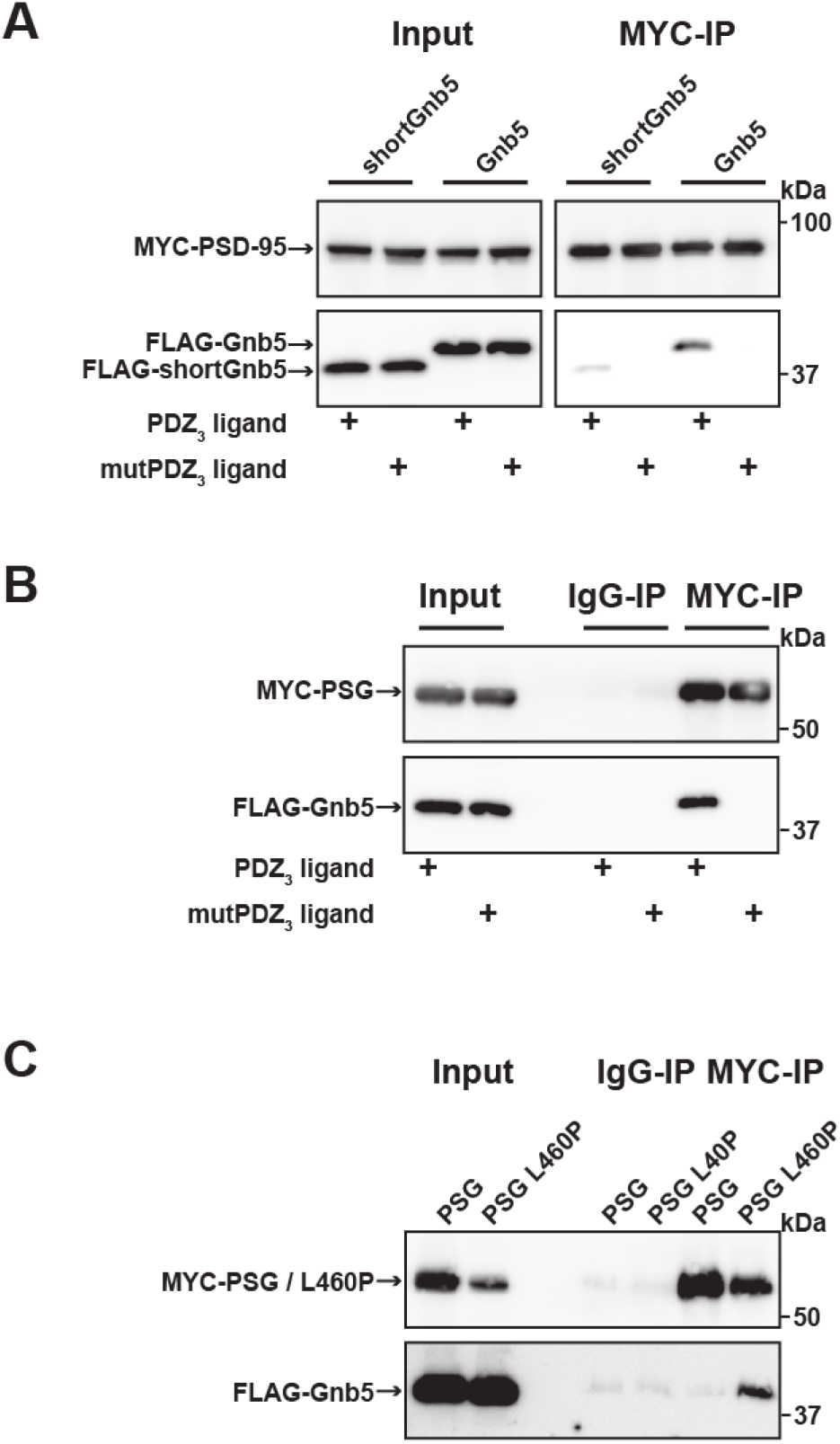
Gnb5 - PSD-95 complex formation is regulated by PDZ_3_ ligand binding. **A)** MYC-PSD-95 and FLAG-Gnb5 or FLAG-shortGnb5 were coexpressed with either PDZ_3_ ligand or mutPDZ_3_ ligand constructs. MYC-PSD-95 was precipitated and proteins were analysed by western blot with αFLAG antibodies. Coexpression of the PDZ_3_ ligand facilitated the coIP of PSD-95 and Gnb5, coIP with the shortGnb5 construct (N-terminal truncation) was much less efficient. In the presence of the mutPDZ_3_ ligand, coprecipitated proteins were not detectable. **B)** CoIP of MYC-PSG and Flag-Gnb5 together with either PDZ_3_ ligand or mutPDZ_3_ ligand constructs. The presence of PDZ3 ligand constructs facilitated coprecipitation of PSG and Gnb5 (see comparative western blot with αFLAG antibodies, lower panel). **C)** Coexpression of MYC-PSG or MYC-PSG L460P with FLAG-Gnb5 and subsequent MYC IP. PSG L460P IP efficiently copurifies Gnb5 (observed by western blot with αFLAG antibodies).

Next, we asked if the PSD-95 PSG module is sufficient to bind to Gnb5 in a ligand-triggered mode. We coexpressed a PSG expression construct together with Gnb5 and PDZ_3_ ligand constructs (wild-type or mutant) and performed pull-downs of the PSG constructs or unspecific IgGs as a control. Upon analysis of the precipitates by western blot, we detected a robust coIP of Gnb5 with the PSG module construct in the presence of PDZ_3_ ligands (**Figure 4B**). Clearly, the PSG module is sufficient for ligand-triggered coimmunoprecipitation of Gnb5.

Our comparative mass spectrometry results for Gnb5, together with subsequent PSD-95 coimmunoprecipitation data, support the idea that ligand binding influences the PSD-95 PSG module such that its protein interaction profile resembles that of the isolated GK domain, *i.e.* it differs from the SH3-GK domain tandem with regard to protein-protein interactions (see **Figure 1D**). In summary, we propose that binding of a PDZ_3_ ligand weakens the intramolecular SH3-GK domain association, which then enables the individual SH3 and GK domains to participate in *trans* interactions with other molecules. To test this model, we took advantage of the L460P mutation, which is known to disrupt the well-characterised intramolecular SH3-GK domain assembly, thus aberrantly releasing the GK domain from its SH3 domain-mediated inhibition. Upon coexpression of wild-type or PSG L460P proteins together with Gnb5, we performed pull-downs of the PSG proteins and comparatively assessed coprecipitation of Gnb5. Gnb5 did not coprecipitate efficiently with the wild-type PSG module but was effectively coprecipitated by the PSG module harbouring the L460P mutation that disrupts the intramolecular SH3-GK domain interaction (**Figure 4C**). We conclude that Gnb5 is interacting with the PSD-95 PSG module in one of two possible modes. Gnb5 could bind at GK domain sites that are directly occupied by the neighbouring SH3 domain (and thereby compete with the SH3 domain for interaction with the GK domain). Alternatively, Gnb5 could bind to GK domain sites on distant surfaces (e.g. the canonical GMP-binding region) that are not directly influenced by intramolecular SH3-GK interactions but might be allosterically regulated by changes to the PSG module.

### PSD-95 interactors occupy different GK subdomains

In order to explore these two possibilities in more depth, we took advantage of established knowledge on the structure of GK domains and information on previously identified GK-interacting proteins. The GK domain of PSD-95 has evolved from an enzyme that catalyses the phosphorylation of GMP to an enzymatically inactive protein interaction domain. Interestingly, various PSD-95 GK-interacting proteins bind to the canonical GMP-binding region, and by exchanging arginine 568 (which is situated in the ancestral GMP-binding site) to alanine (R568A), these interactions can be specifically disrupted (Reese et al., 2007). In order to gain insight into the nature of the binding of Gnb5 to the PSD-95 GK domain, we compared PSD-95 - Gnb5 binding to PSD-95 - GKAP binding. GKAP (‘GK’-associated protein, also referred to as SAPAP1 or DLGAP1) is an established synaptic GK domain binder (Kim et al., 1997) whose interaction involves the GMP-binding region (Zhu et al., 2017). These ideas are also validated by our own coimmunoprecipitation experiments: GKAP can be efficiently coprecipitated upon pull-down of either the isolated GK domain or an intact PSG module, whereas a recombinant PSG module harbouring the GMP binding site mutation R568A fails to precipitate GKAP (**Figure 5A**). In experiments with PSD-95 and Gnb5, however, the same mutation had no effect on coprecipitation of Gnb5 (**Figure 5B**), suggesting that GKAP and Gnb5 proteins bind to PSD-95 GK domains in fundamentally different ways.

**Figure 5.**
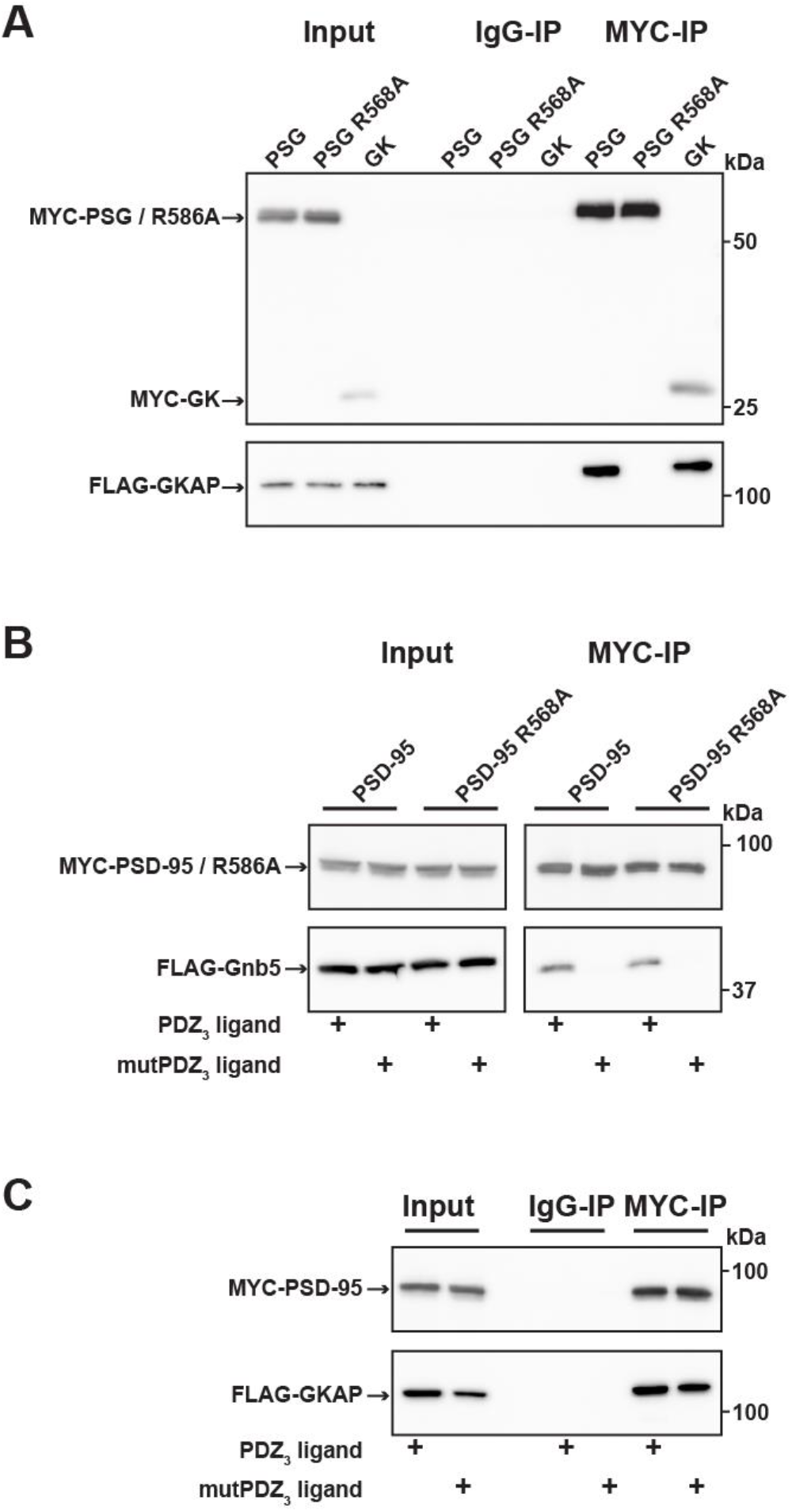
PSD-95 interactors occupy different protein surfaces. **A)** MYC-PSG, MYC-PSG R568A and MYC-GK were coexpressed with FLAG-GKAP. Following MYC IP, precipitated proteins were analysed by western blot. GKAP coprecipitated with PSG and GK domain constructs. The GK domain mutant PSG R568A was not able to bind GKAP. **B)** CoIP of MYC-PSD-95 and FLAG-GKAP together with either PDZ_3_ ligand or mutPDZ_3_ ligand constructs and analysis of precipitated proteins by western blot with antibodies to the corresponding tags. The presence of PDZ_3_ ligands in the lysate had no effect on PSD-95 GKAP interaction. **C)** Following coexpression of MYC-PSD-95 or MYC-PSD-95 R568A with FLAG-Gnb5, together with either PDZ_3_ ligand or mutPDZ_3_ ligand, proteins were precipitated with αMYC-antibody and analysed by western blot. Gnb5 coIP with either PSD-95 or PSD-95 R568A was efficiently promoted by the presence of PDZ_3_-binding ligand, irrespective of the GK domain mutation R568A.

We next tested whether the GKAP - PSD-95 association could be influenced by PDZ_3_ ligands that bind to PSD-95, as we observed previously for Gnb5 (see **Figure 4A, 4B and 5B**). The presence of PDZ_3_ ligands did not influence the GKAP interaction: PSD-95 binds GKAP regardless of whether wild-type or mutant PDZ_3_ ligands were present (**Figure 5C**). These data provide further evidence that the GKAP - GK domain binding mode differs substantially from the Gnb5 - GK interaction mode.

Importantly, our data support a model in which ligand binding to PDZ_3_ results in a conformational change of the ‘resting’ intramolecular SH3-GK interaction that is common to MAGUK proteins. This conformational alteration is reflected by a change in the availability of specific GK surfaces for protein-protein interactions (**Figure 6A**). In the resting state, it is predominantly the external GK surface harbouring the classical GMP binding site that is available for protein-protein interaction such as those with the well-known PSD-95 interactors GKAP and MAP1a. However, upon ligand binding, other GK surfaces become accessible for protein-protein interactions. A subset of synaptic GK interacting proteins – in particular Gnb5, and perhaps other proteins enriched in our pool of interacting proteins that bind preferentially to GK rather than to SH3-GK – bind to these surfaces of the GK domain (**Figure 6B**).

**Figure 6.**
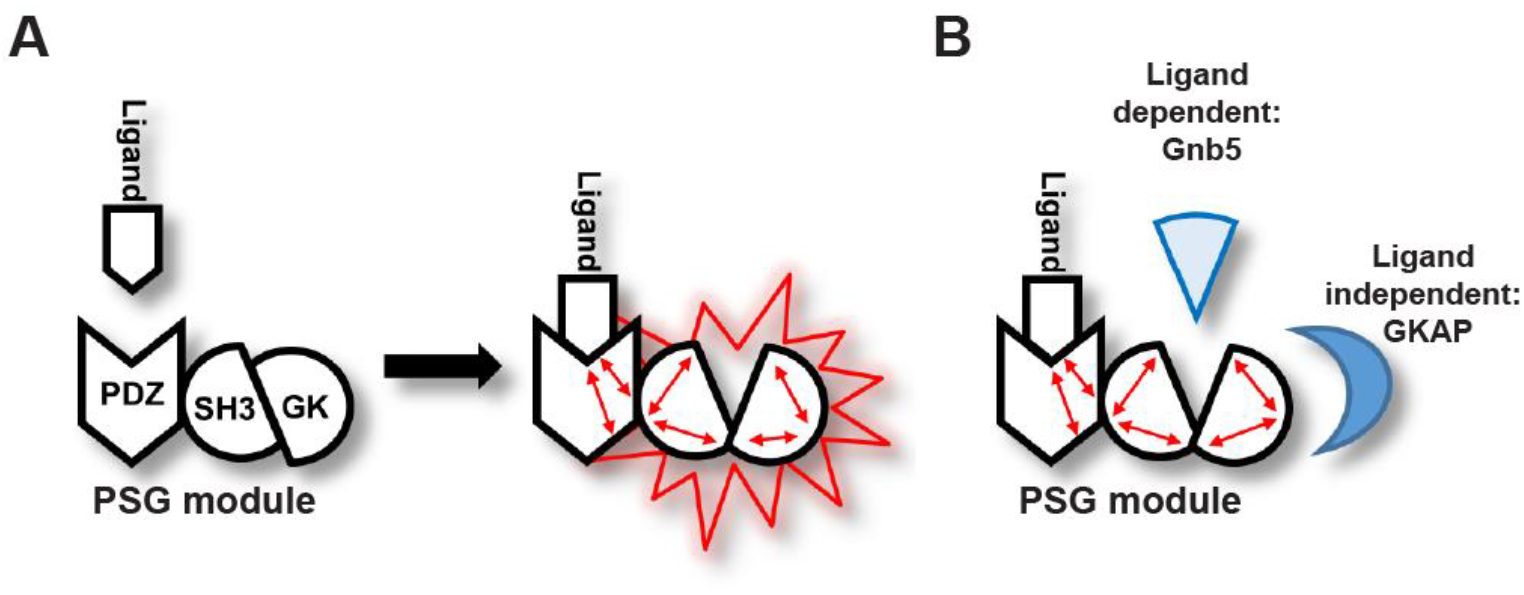
Graphical Summary. **A)** PSD-95 C-terminal domains (PSG module) functionally cooperate and regulate homotypic and heterotypic complex formation. We propose that PDZ_3_ ligand binding to the PDZ_3_ domain induces a loosening of the intramolecular SH3-GK interaction. This ‘open’ conformation is then able to initiate subsequent oligomerisation and protein binding. **B)** Model of PDZ_3_ ligand-dependent and ligand-independent binding to the PSD-95 C terminal SH3-GK domain tandem. Ligand - PDZ_3_ domain binding facilitates association with Gnb5.

### Discussion

The molecular basis for the dynamic regulation of synaptic transmission is dependent on the assembly and disassembly of protein complexes. It has been observed that activation of glutamate receptors is sufficient to specifically remodel postsynaptic protein-protein interactions (Lautz et al., 2018). In line with this finding is that upon LTP induction, various proteins undergo post-translational modifications and are incorporated into dense protein complexes at the postsynaptic compartment (Yokoi et al., 2012). It is well established that these activity-dependent changes in synaptic protein networks depend on phosphorylation (Opazo et al., 2010; Araki et al., 2015; Li et al., 2016) and other post-translational modifications, such as palmitoylation (El-Husseini Ael et al., 2002; Fukata et al., 2013). Recently it has been reported that the minimal requirement for the expression of LTP is the interaction of glutamate receptor auxiliary subunits with postsynaptic PSD-95. In that study, the interaction of different PDZ ligand C-termini with PSD-95 triggered a common molecular mechanism necessary for LTP induction downstream of glutamate receptors (Sheng et al., 2018).

In this study, we focussed on the postsynaptic scaffold protein PSD-95, which plays a central role in activity-dependent synapse regulation (Ehrlich et al., 2007). It is established that protein complex formation guided by PSD-95 PDZ and GK domains can be reversibly regulated by phosphorylation (Sumioka et al., 2010; Zhu et al., 2017), and at postsynaptic membranes, various PDZ ligand C-termini of multimeric receptor complexes are available to form multivalent interactions with scaffold proteins (Schwenk et al., 2012). Importantly, in a previous study it was also shown that binding of a SynGAP-derived PDZ ligand peptide was sufficient to induce PSD-95 PSG construct dimerisation (Zeng et al., 2016), but the underlying mechanism remained unresolved. Here, we show that PSD-95 oligomerisation can be induced by binding of monomeric PDZ_3_ ligands, which then leads to conformational changes in the adjacent C-terminal SH3-GK domain structure.

Moreover, we identify synaptic interactors whose association with PSD-95 is likewise influenced by the conformational state of the PSD-95 C-terminus. Among these proteins, we focussed further on Gnb5, which is part of a protein complex that acts downstream of GABA_B_ receptors and also modulates GIRK channel gating properties (Ostrovskaya et al., 2014). Our data indicate that the Gnb5 - PSD-95 interaction is positively regulated by ligand binding to the third PDZ domain of PSD-95. In order to understand how this occurs, it is important to note that in MAGUK scaffold proteins, the SH3 and GK domains interact directly, and together they form a unique structure that sets them apart from SH3 and GK domains found independently in other protein families (Tavares et al., 2001). Indeed, PSD-95 SH3 and GK domains, when expressed independently, bind each other efficiently. Likewise, in line with this structural model, mutations that disrupt the interface where these two domains contact each other can have detrimental effects on protein function (McGee and Bredt, 1999; Shin et al., 2000). Also relevant is the fact that the SH3 domain has been reported to be an allosteric regulator or inhibitor of GK domain binding function, not only by direct contact with the adjacent GK surface but also by regulating the conformation of distant GK domain surfaces (Marcette et al., 2009). It is possible that binding of a ligand to the PSD-95 PDZ_3_ domain influences precisely this function of the neighbouring SH3 domain and thus indirectly regulates GK interactions at distant sites. Alternatively, it is conceivable that regulation via PDZ_3_ ligand binding results in a conformational change that loosens the natural SH3-GK structure, thereby freeing up the SH3-interacting surface of the GK domain for other protein-protein interactions. Nevertheless, in both possible scenarios, binding of a ligand to the adjacent PDZ_3_ domain would release the GK domain from its regulation by the interacting SH3 domain. In order to explore these two possibilities, we took advantage of the established GK interactor GKAP and we compared Gnb5 and GKAP with regard to PSD-95 binding. By introducing a mutation (R568A) in the canonical GMP-binding region of PSD-95, we were able to completely abolish GKAP binding to the PSD-95 GK domain. The GKAP - PSD-95 interaction, however, was not influenced by ligand binding to PDZ_3_. This result suggests that the canonical GMP-binding region in the GK domain is *not* allosterically regulated by PDZ_3_ ligand binding. For Gnb5 we observed the opposite pattern: First of all, the R568A mutation had no effect on the Gnb5 - PSD-95 interaction, enabling us to conclude that Gnb5 occupies different GK domain surfaces for interaction, indicating that Gnb5 occupies different GK domain surfaces for interaction that do not overlap with the canonical GMP-binding site responsible for the GKAP interaction. Second, the Gnb5 - PSD-95 association, unlike the GKAP - PSD-95 interaction, is strongly dependent on ligand binding to PDZ_3_, further corroborating the idea that Gnb5 binds the PSD-95 GK domain away from the GMP-binding site.

We propose that the Gnb5 - PSD-95 interaction is regulated by a modular allosteric mechanism: the SH3 domain exerts inhibitory activity on the GK domain binding capacity by competing directly with Gnb5 for interaction surfaces. The PSD-95 SH3-GK domain tandem undergoes structural rearrangements upon binding of a PDZ_3_ ligand to the adjacent PDZ_3_ domain, and these changes free up the GK domain for interactions with selected proteins. Via this mechanism, ligand - PDZ_3_ domain interactions facilitate formation of both homotypic and heterotypic complexes guided by the PSD-95 C-terminal PSG domain module.

## Materials and Methods

### DNA Constructs

Full-length rat PSD-95 (NM_019621) was cloned into pCMV-Tag3A, to obtain MYC-PSD-95. Arginine 568 was exchanged to Alanine by site-directed mutagenesis to generate MYC-PSD-95 R568A.

PSD-95-YN was generated by a PCR based strategy: amino acids 1 − 723 of PSD-95 were fused to a flexible 3x(GGGGS) linker followed by amino acids 1 − 154 of EYFP and an HA-tag. PSD-95-YC was generated accordingly by fusing amino acids 1 − 723 of PSD-95 to a flexible 3x(GGGGS) linker followed by amino acids 155 − 238 of EYFP and a MYC-tag. Leucine 460 was exchanged to Proline by site-directed mutagenesis to generate PSD-95-YN L460P.

MYC-PSG was generated by cloning a fragment that encodes amino acids 247 – 724 of PSD-95 into pCMV-Tag3A. MYC-PSG L460P and MYC-PSG R568A were generated by PCR based site-directed mutagenesis. FLAG-SH3-GK and FLAG-GK constructs were generated by cloning fragments that encode amino acids 403 – 724 (SH3-GK) and amino acids 504 − 724 (GK) of PSD-95 into pCMV-Tag2A. MYC-GK was generated by cloning a fragment that encodes amino acids 504 − 724 of PSD-95 into pCMV-Tag3A.

GST-SH3-GK and GST-GK constructs were generated by cloning fragments that encode amino acids 403 − 724 (SH3-GK) and amino acids 504 − 724 (GK) of PSD-95 into pGEX-6P-1 (GE Healthcare).

Full-length rat Neuroligin-1 (NLGN1, NM_053868.2) was cloned into pcDNA3.1 and an HA-tag (YPYDVPDYA) was inserted following the signal sequence (between amino acid 45 and 46) by PCR mutagenesis. The C-terminal PDZ ligand motif (TTRV) was mutated to abolish PDZ domain binding by introducing two alanine residues (T**A**R**A**) to generate a mutNLGN1 construct.

PDZ ligand constructs were generated by fusing an HSV-tag (QPELAPEDPED) to mCherry followed by a flexible 3x(GGGGS) linker and 10 aminoacids (DTKNYKQTSV) referring to the PDZ ligand CRIPT. MutPDZ ligand constructs were generated by mutating the C-terminal QTSV motif to Q**A**S**A.**

Full-length rat Gnb5 (NM_031770) was cloned into pCMV-Tag2A to generate FLAG-Gnb5. A shortGnb5 construct was generated by cloning a fragment that encodes amino acids 34 – 353 of Gnb5 into pCMV-Tag2A. Gnb5-Clover was generated by cloning Gnb5 into pEYFP-N1. In a subsequent clonig step EYFP was exchanged for Clover. Full-length mouse Gkap (NM_001360665) was cloned into pCMV-Tag2A to generate FLAG-mGkap.

### Cell Culture and Transfection

COS-7 and HEK-293T cells were maintained in DMEM containing 10% FCS, PEN-STREP (1,000 U/ml) and 2 mM L-glutamine. Cells were transfected with Lipofectamine 2000 Reagent (Invitrogen) according to the manufacturer’s protocol and harvested for subsequent experimental procedures 20–24 hours post transfection.

### Bimolecular fluorescence complementation (BiFC) assay and flow cytometry

HEK-293T cells were cultured in 12 well plates and transfected with the respective expression construct combinations. Prior to analysis by flow cytometry (BD FACS Calibur) the cells were incubated for 60 minutes at room temperature to promote fluorophore formation. Cells were harvested by gently washing the culture dishes with PBS / 10% FCS. 10,000 single-cell events for each construct combination were measured and fluorescence was quantified (BD CellQuest).

### Coimmunoprecipitation

Transfected COS-7 cells were harvested 20–24 hours post transfection, resuspended in lysis buffer (50 mM Tris-HCl, 100 mM NaCl, 0.1% NP40, pH 7.5 / 1 ml per T75 flask) and lysed using a 30-gauge syringe needle. Lysates (1 ml) were cleared by centrifugation and incubated with 2 mg of the appropriate antibody (mouse αGFP antibody [Roche], mouse αMYC [Clontech], or normal mouse IgG [Santa Cruz]) for 3 hours followed by a centrifugation at 20,000 × g. Supernatants were incubated with 30 μl Protein G-Agarose (Roche) per ml and washed three times with lysis buffer.

### Western Blot

Immunocomplexes were collected by centrifugation, boiled in SDS sample buffer, and separated by 10% Tricine-SDS-PAGE (Schagger, 2006). Proteins were blotted onto a PVDF membrane (0.2 mm pore size, Bio-Rad) by semidry transfer (SEMI-DRY TRANSFER CELL, Bio-Rad). Membranes were blocked (PBS / 0.1% Tween 20 / 5% dry milk) and incubated overnight with the primary antibody (1:5000). After incubation with the respective horseradish peroxidase (HRP)-conjugated secondary antibody (1:5000), blots were imaged using chemiluminescence HRP substrate (Western Lightning Plus ECL, Perkin Elmer) and a luminescent image analyzer (ImageQuant LAS 4000 mini, GE Healthcare). The following primary antibodies were used for protein detection: αFLAG M2-HRP (mouse, A8592, Sigma), αGnb5 (rabbit, ab185206, Abcam), αMYC (rabbit, 2272S, Cell Signalling). Secondary antibodies: αMouse-HRP (115-035-003, Dianova), αRabbit-HRP (111-035-003, Dianova). All western blots shown are representative results from individual pull-down experiments that have been replicated at least three times with similar outcome.

### Isolation of crude synaptosomes and GST pull-down

One rat brain (Wistar, 2g) was used to isolate synaptic proteins with Syn-PER reagent (Thermo Scientific) according to the manufacturer’s manual. The purified synaptosome pellet was solubilised in 10ml PBS / 1% Triton X-100 and cleared by centrifugation.

GST-GK and GST-SH3-GK constructs were expressed in *E.coli* BL21 DE3 and purified according to the manufacturer’s manual (GST Gene Fusion System, GE Healthcare). 30μl of Glutathione Agarose (Pierce) was loaded with GST-GK or GST-SH3-GK proteins and incubated for 3 hours with solubilised synaptic proteins. The beads were washed three times with PBS / 1% Triton X-100 and further processed for SDS-PAGE.

### Sample preparation and liquid chromatography-mass spectrometry (LC-MS)

Proteins were eluted from the matrix by incubation with SDS sample buffer for 5 min at 95 °C and subsequently separated by SDS-PAGE (10% Tricine-SDS-PAGE). Coomassie-stained lanes were cut into 12 slices and in-gel protein digestion and ^16^O/^18^O-labeling was performed as described (Kristiansen et al., 2008; Lange et al., 2010). In brief, corresponding samples were incubated overnight at 37 °C with 50 ng trypsin (sequencing grade modified, Promega) in 25 μL of 50 mM ammonium bicarbonate in the presence of heavy water (Campro Scientific GmbH, 97% ^18^O) and regular ^16^O-water, respectively. To prevent oxygen back-exchange by residual trypsin activity, samples were heated at 95 °C for 20 min. After cooling down, 50 μL of 0.5% TFA in acetonitrile was added to decrease the pH of the sample from ~8 to ~2. Afterwards, corresponding heavy- and light-isotope samples were combined and peptides were dried under vacuum. Peptides were reconstituted in 10 μL of 0.05% (v/v) TFA, 2% (v/v) acetonitrile and 6.5 μL were analyzed by a reversed-phase capillary nano liquid chromatography system (Ultimate 3000, Thermo Scientific) connected to an Orbitrap Velos mass spectrometer (Thermo Scientific). Samples were injected and concentrated on a trap column (PepMap100 C18, 3 μm, 100 Å, 75 μm i.d. × 2 cm, Thermo Scientific) equilibrated with 0.05% TFA, 2% acetonitrile in water. After switching the trap column inline, LC separations were performed on a capillary column (Acclaim PepMap100 C18, 2 μm, 100 Å, 75 μm i.d. × 25 cm, Thermo Scientific) at an eluent flow rate of 300 nL/min. Mobile phase A contained 0.1% formic acid in water, and mobile phase B contained 0.1% formic acid in acetonitrile. The column was pre-equilibrated with 3 % mobile phase B followed by an increase of 3–50% mobile phase B in 50 min. Mass spectra were acquired in a data-dependent mode utilizing a single MS survey scan (m/z 350-1500) with a resolution of 60,000 in the Orbitrap, and MS/MS scans of the 20 most intense precursor ions in the linear trap quadrupole. The dynamic exclusion time was set to 60 s and automatic gain control was set to 1 × 106 and 5.000 for Orbitrap-MS and LTQ-MS/MS scans, respectively.

### Proteomic Data Analysis

Identification and quantification of ^16^O/^18^O-labeled samples was performed using the Mascot Distiller Quantitation Toolbox (version 2.6.3.0, Matrix Science). Data were compared to the SwissProt protein database using the taxonomy *rattus* (August 2017 release with 7996 protein sequences). A maximum of two missed cleavages was allowed and the mass tolerance of precursor and sequence ions was set to 10 ppm and 0.35 Da, respectively. Methionine oxidation, acetylation (protein N-terminus), propionamide (C), and C-terminal ^18^O_1_- and ^18^O_2_- isotope labeling were used as variable modifications. A significance threshold of 0.05 was used based on decoy database searches. For quantification at protein level, a minimum of two quantified peptides was set as a threshold. Relative protein ratios were calculated from the intensity-weighted average of all peptide ratios. The median protein ratio of each experiment was used for normalization of protein ratios. Only proteins that were quantified in all three replicates with a standard deviation of < 2 were considered. Mean protein L/H ratios (GST-GK /GST-SH3-GK) from all three replicates were calculated. Known contaminants (e.g. keratins) and the bait protein were removed from the protein output table.

### Live cell microscopy

HEK-293T cells were seeded in 35 mm FluoroDishes (World Precision Instruments) and triple-transfected with PSD-95-YN, PSD-95-YC and Neuroligin-1 expression constructs. Images were acquired using a spinning disk confocal microscope (Nikon CSU-X).

### Immunofluorescence and confocal microscopy

Mixed cultures of primary hippocampal neurons were generated as reported earlier (Rademacher et al., 2016). Briefly, E18 Wistar pups were decapitated, and hippocampi were isolated and collected in ice-cold DMEM (Lonza). Single cell solution was generated by partially digestion (5 min at 37 °C) with Trypsin/EDTA (Lonza). The reaction was stopped by adding DMEM/10% FBS (Biochrom) following a subsequent washing with DMEM. Tissue was then resuspended in neuron culture medium (Neurobasal supplemented with B27 and 500 μ M glutamine) and mechanically dissociated. Neurons were plated at ~105 cells/cm2 on coverslips coated with poly-D-Lysine and Laminin (Sigma). One hour after plating, cell debris was removed and cultures were maintained in a humidified incubator at 37 °C with 5% CO2. The hippocampal neurons were fixed at DIV21 with 4% PFA in PBS for 10 min at RT and permeabilised with 0.2% Triton-X in PBS for 5 min. After blocking for 1 h at RT with blocking solution (4% BSA in PBS) the primary antibodies were incubated overnight at 4°C diluted 1:500 in blocking solution. Secondary antibodies were diluted 1:1000 in blocking solution and incubated for 1 hour at RT. Coverslips were mounted with Fluoromount G and images were acquired with a Leica laser-scanning confocal microscope (Leica TCS-SP5 II, 63x objective). Primary antibodies: αGnb5 (rabbit, ab185206, Abcam), αPSD-95 (mouse, 75-028, NeuroMab), αMAP2 (mouse, 05-346, Millipore), αMAP2 (guinea pig, 188004, Synaptic Systems), αSynapsin (guinea pig, 106004, Synaptic Systems). Secondary antibodies: αRabbit Alexa Fluor 488 (A-21441, Invitrogen), αGuinea pig Alexa Fluor 568 (Thermo Fisher), αGuinea pig Alexa Fluor 405 (ab175678, Abcam), αMouse Alexa Fluor 568 (A-11031, Life Technologies), αMouse Alexa Fluor 405 (A-31553, Invitrogen).

### Laboratory animal handling

All animals used were handled in accordance with the relevant guidelines and regulations. Protocols were approved by the ‘Landesamt für Gesundheit und Soziales’ (LaGeSo; Regional Office for Health and Social Affairs) in Berlin and animals reported under the permit number T0280/10.

## Author Contributions

N.R., B.K. and S.-A.K. performed experiments. N.R., C.F., M.C.W. and S.A.S. designed experiments and analysed data, N.R. and S.A.S. wrote the paper.

## Acknowledgments

We would like to thank Bettina Schmerl and Hanna Zieger for helpful comments and support, the Advanced Medical Bioimaging Core Facility – Excellence Center for Microscopy at the Charité Berlin for support with spinning disk microscopy imaging. This work was funded by the DFG (SH 650/2, Collaborative Research Centres SFB 958 / SFB 665 and Excellence Cluster NeuroCure EXC257).

## Competing interests

The authors declare no competing interests.

---

**Supplemental Figure 1.**
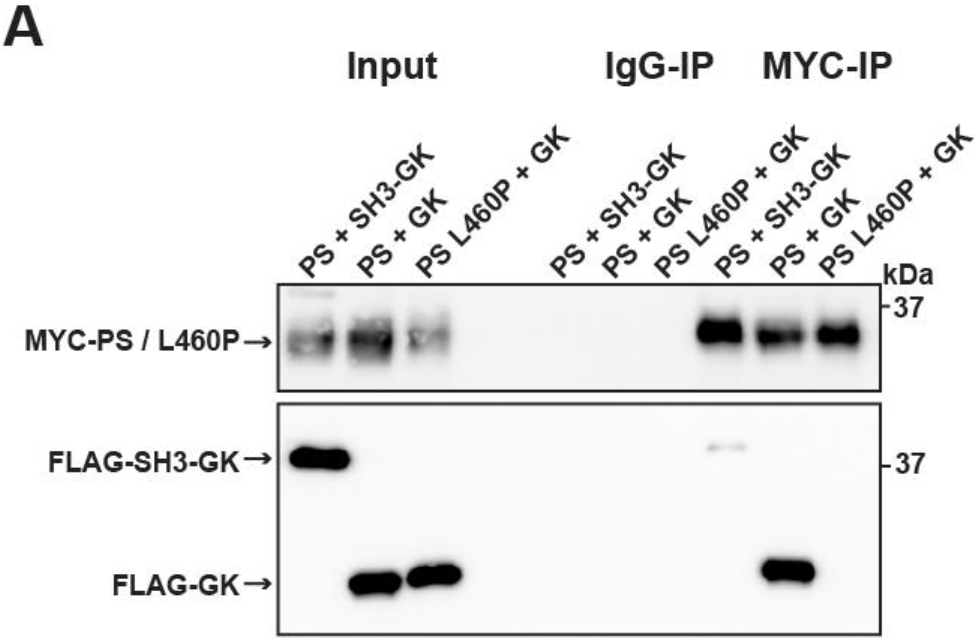
**A)** PSD-95 constructs consisting of the PDZ3-SH3 domains (PS) were coexpressed together with either an SH3-GK domain construct, or a GK domain costruct. As a comparison PDZ3-SH3 L460P was coexpressed with a GK domain construct and PDZ3 SH3 / PDZ3-SH3 L460P constructs were precipitated and copurified proteins were identified by western blot. By mutating the leucine 460 to proline this efficient protein complex formation is disrupted. By exchanging the internal L460 residue the SH3 domain loses its ability to bind to the GK domain construct in trans.

